# Expanding Asgard members in the domain of Archaea shed new light on the origin of eukaryotes

**DOI:** 10.1101/2021.04.02.438162

**Authors:** Ruize Xie, Yinzhao Wang, Danyue Huang, Jialin Hou, Liuyang Li, Haining Hu, Xiaoxiao Zhao, Fengping Wang

## Abstract

The hypothesis that eukaryotes originated from within the domain Archaea has been strongly supported by recent phylogenomic analyses placing Heimdallarchaeota from the Asgard superphylum as the closest known archaeal sister-group to eukaryotes. At present, only six phyla are described in the Asgard superphylum, which limits our understanding of the relationship between eukaryotes and archaea, as well as the evolution and ecological functions of the Asgard archaea. Here, we describe five previously unknown phylum-level Asgard archaeal lineages, tentatively named Tyr-, Sigyn-, Freyr-, Njord- and Balderarchaeota. Comprehensive phylogenomic analyses further supported the origin of eukaryotes within Archaea and a new Asgard lineage Njordarchaeota was supposed as the known closest branch with the eukaryotic nuclear host lineage. Metabolic reconstruction suggests that the Asgard archaea described here have potential to fix inorganic carbon via the Wood-Ljungdahl pathway and degrade organic matters except Njordarchaeota, which may possess a heterotrophic lifestyle with capability of peptides and amino acids utilization. Additionally, the Ack/Pta pathway for homoacetogenesis and *de novo* anaerobic cobalamin biosynthesis pathway were found in Balderarchaeota and Tyrarchaeota, respectively. This study largely expands the Asgard superphylum, provides additional evidences to support the 2-domain life tree and sheds new light on the evolution of eukaryotes.

## Introduction

The origin of eukaryotes is considered as a critical biological evolutionary event on Earth (Embley and Martin 2006, López-García and Moreira 2015, Rochette et al., 2014, Lopez-García and Moreira 2006). The common ancestor of eukaryotes is generally believed to have evolved from a symbiotic process (López-Garcia and Moreira 1999, Martin et al., 2015) in which one endosymbiotic bacterium within the Proteobacteria phylum evolved into a mitochondrion (Esser et al., 2004, Moreira and López-García 1998) and one endosymbiotic host cell became the cell nucleus (Spang et al., 2015, Zaremba-Niedzwiedzka et al., 2017a, Spang et al., 2019a). The identity of the host cell ancestor has been vigorously debated, and two hypotheses regarding 2- or 3-domain trees of life have been raised (Williams et al., 2013, Woese et al., 1990). However, increasing evidence provided by phylogenomic analyses (Zaremba-Niedzwiedzka et al., 2017a, Williams et al., 2020), as well as the presence of eukaryotic signature proteins (ESPs) (Hartman and Fedorov 2002) in the Asgard archaea, has supported the idea that eukaryotic cells originated from an archaeal host cell related to archaeal Asgard superphylum (Spang et al., 2015, Zaremba-Niedzwiedzka et al., 2017a, Bulzu et al., 2019). The identification of Lokiarchaeota in the Loki’s Castle hydrothermal vent field provided pivotal genomic and phylogenetic evidence that eukaryotes originated within the domain Archaea, supporting a 2-domain tree of life, which is consistent with the eocyte hypothesis (Spang et al., 2015). Further discovery and proposal of the Asgard superphylum have provided new insights into the transition of archaea to eukaryotes and into the origin of eukaryotic cell complexity (Zaremba-Niedzwiedzka et al., 2017a). Within the Asgard superphylum, Heimdallarchaeota had been identified to be the closest Asgard archaeal lineage to the eukaryotic branch on the phylogenetic tree on the basis of carefully selected conserved protein sequences (Zaremba-Niedzwiedzka et al., 2017a, Williams et al., 2020). Recently, Imachi *et al.* cultivated one Asgard archaeon, *Candidatus* Prometheoarchaeum syntrophicum strain MK-D1, in the laboratory and observed, for the first time, the intertwining of this archaeon with bacterial cells via extracellular protrusions under a transmission electron microscope (Imachi et al., 2020). The idea of an archaeal origin of eukaryotes and a 2-domain tree of life has recently become increasingly favorable (Williams et al., 2020, Akil et al., 2020). The Asgard archaea are described as mixotrophic or heterotrophic (Spang et al., 2019a, Liu et al., 2018a) and are ubiquitously distributed in various environments, such as hydrothermal vents (Spang et al., 2015, Zaremba-Niedzwiedzka et al., 2017a); lake, river and marine sediments (Seitz et al., 2016); microbial mats (Wong et al., 2018); and mangroves (Liu et al., 2018a). These organisms potentially play important roles in global geochemical cycling (MacLeod et al., 2019); nevertheless, our understanding of the evolution of Asgard archaea, the archaea-eukaryote transition, and the ecological and geochemical roles of these evolutionarily important archaea remains incomplete. This lack of understanding is largely due to the limited number of high-quality genomes of Asgard archaea, which are considered highly diverse as revealed by 16S rRNA gene surveys (MacLeod et al., 2019, Zhang et al., 2020); yet only a small fraction have representative genomes. In this study, we described five previously unknown phylum-level Asgard archaeal group, greatly expanding the Asgard genomic diversity within the domain of Archaea and shed new light on the origin of eukaryotes.

## Results and Discussion

### Expanded Asgard archaea support 2-domain tree of life

In total, 11 metagenomic datasets were used in this study, including two samples from hydrothermal sediment of Guaymas Basin, six samples from Tengchong hot spring sediment, as well as 3 metagenomic datasets from the publicly available National Center for Biotechnology Information (NCBI) Sequence Read Archive (SRA) database (Permission granted, Supplementary Table 1). After subsequent assembling, binning and classification as described in the Methods section, 128 Asgard metagenome-assembled genomes (MAGs) were obtained and in-depth phylogenomic analyses were performed with 37 concatenated conserved proteins (Wang et al., 2019) under LG+C60+F+G4 model to confirm the placement of these MAGs on phylogenomic tree. The analysis revealed that, in addition to the previously described Loki-, Thor-, Odin-, Heimdall-, Hela- and Hermodarchaeota clades (Zaremba-Niedzwiedzka et al., 2017b, Zhang et al., 2021), there are five additional monophyletic branching clades (Fig. 1a), here tentatively named Tyr-, Sigyn-, Freyr-, Njord- and Balderarchaeota after the Asgard gods in the Norse mythology (Tyr, the god of war; Sigyn, the god of victory; Freyr, the god of peace; Njord, the god of seas; and Balder, the god of light). The MAGs of these new Asgard lineages were recovered from different environments: Njordarchaeota and Freyrarchaeota were derived from hydrothermal sediment; Tyrarchaeota were found in estuary sediments; Sigynarchaeota were reconstructed from hot spring sediments and Balderarchaeota were retrieved from hot spring and hydrothermal sediments. Additionally, a clade of Hermodarchaeota was identified in high temperature habitats (~85°C), similar with Odinarchaeota, which were considered as the only thermophilic member of Asgard archaea to date (Zaremba-Niedzwiedzka et al., 2017a). Near-complete MAGs ranging in size from 2.1 to 5.5 Mb with completeness ranging from 87.38 to 97.20% were constructed for representatives of each new Asgard clade (Supplementary Table 2). To further assess their distinctiveness compared to the Asgard members already defined, we calculated the average nucleotide identity (ANI) (Supplementary Fig. 1) and average amino acid identity (AAI) (Supplementary Fig. 2) between them and other Asgard MAGs. The AAI values showed that all the MAGs of new lineages discovered here share a low AAI with the known Asgard archaea (<50%) and fall within the phylum-level classification range (40%~52%) (Luo et al., 2014), providing additional support for the uniqueness of these new Asgard lineages.

**Figure 1.**
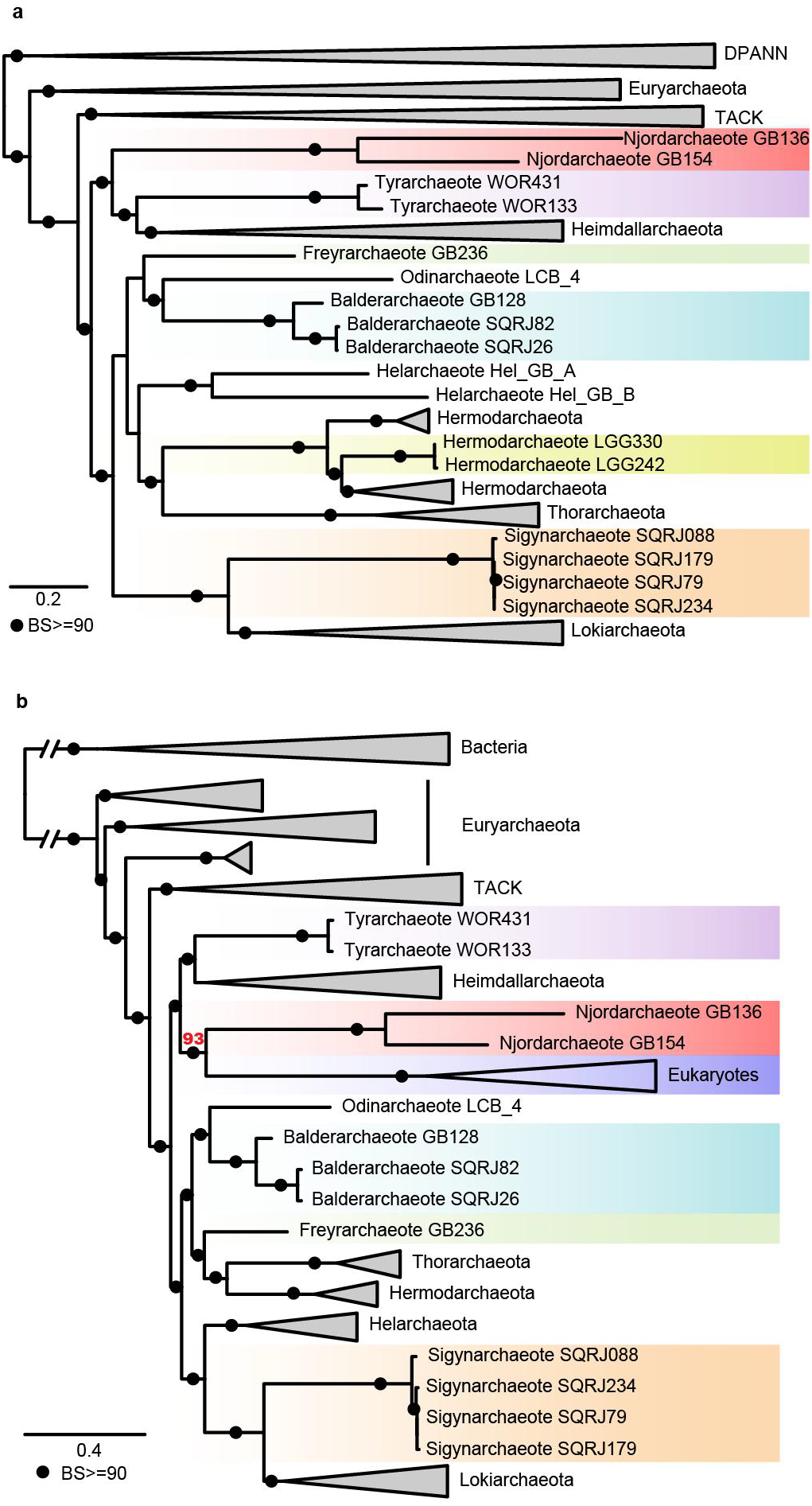
Phylogenetic analyses of Asgard archaea. **a,** Maximum-likelihood tree of 37 concatenated marker proteins inferred with the LG+F+C60+G4 model in IQ-TREE rooted in DPANN, 19 representatives of DPANN, 55 representatives of Euryarchaeota, 29 representatives of TACK and 68 genomes (including five new lineages) of Asgard were used to infer phylogenetic tree. **b**, Maximum likelihood inference of 21 concatenated conserved protein sequences under the LG+F+C60+G4 model rooted in bacteria, 85 archaeal genomes (53 within Asgard), 19 eukaryotic genomes, and 36 bacterial genomes were used to infer phylogenetic tree. The bootstrap support values above 90 were shown with black filled circles.

To determine the phylogenetic positions of these new Asgard lineages in relation to eukaryotes, we performed comprehensive phylogenetic analyses using 21 conserved marker proteins carefully selected by Williams *et al*. (Williams et al., 2020). The taxa included in these analyses were also selected on the basis of the instructions of Williams *et al.* (Williams et al., 2020): a representative taxon set was constructed comprising 85 archaeal genomes (53 within Asgard), 19 eukaryotic genomes, and 36 bacterial genomes (Supplementary Table 3). Single-gene datasets were also carefully inspected with single protein trees to avoid potential phylogenetic artifacts resulting from HGT, long branch attraction (LBA) and BLASTp inspection was performed to remove eukaryotic genes from the mitochondria or plastids. After concatenating, the maximum likelihood tree under the LG+C60+F+G4 model for 21 conserved marker proteins was inferred. The phylogenetic analysis of 21 marker genes suggested that eukaryotes are the sister group of Njordarchaeota with high support (bootstrap support (BS) = 93, Fig. 1b). The phylogenomic analyses provide strong support for a 2-domain tree with Njordarchaeota as the closest potential relatives to eukaryotes.

### ESP-encoding genes widely shared by the Asgard archaea

The potential ESPs were identified from the newly reconstructed Asgard MAGs (Fig. 2). Consistent with previous reports (Spang et al., 2015, Zaremba-Niedzwiedzka et al., 2017a, Bulzu et al., 2019, Seitz et al., 2019), different key subunits of informational processing machinery were found. For example, topoisomerase IB protein-encoding genes were identified in Balderarchaeota and Hermodarchaeota, while all MAGs of Balderarchaeota were found to encode a RNA polymerase subunit G. Homologues of eukaryotic ribosomal protein L22e were identified in Freyr-, Hermodarchaeota and Tyrarchaeota. The new clades of Asgard archaea were also found to contain genes related to cell division and the cytoskeleton, but tubulin-encoding genes were not detected. With regard to actin-related proteins, two to three related subunits were detected in Tyr- and Balderarchaeota, whereas profilin domain protein-encoding genes were identified in all Asgard lineages described here.

**Figure 2.**
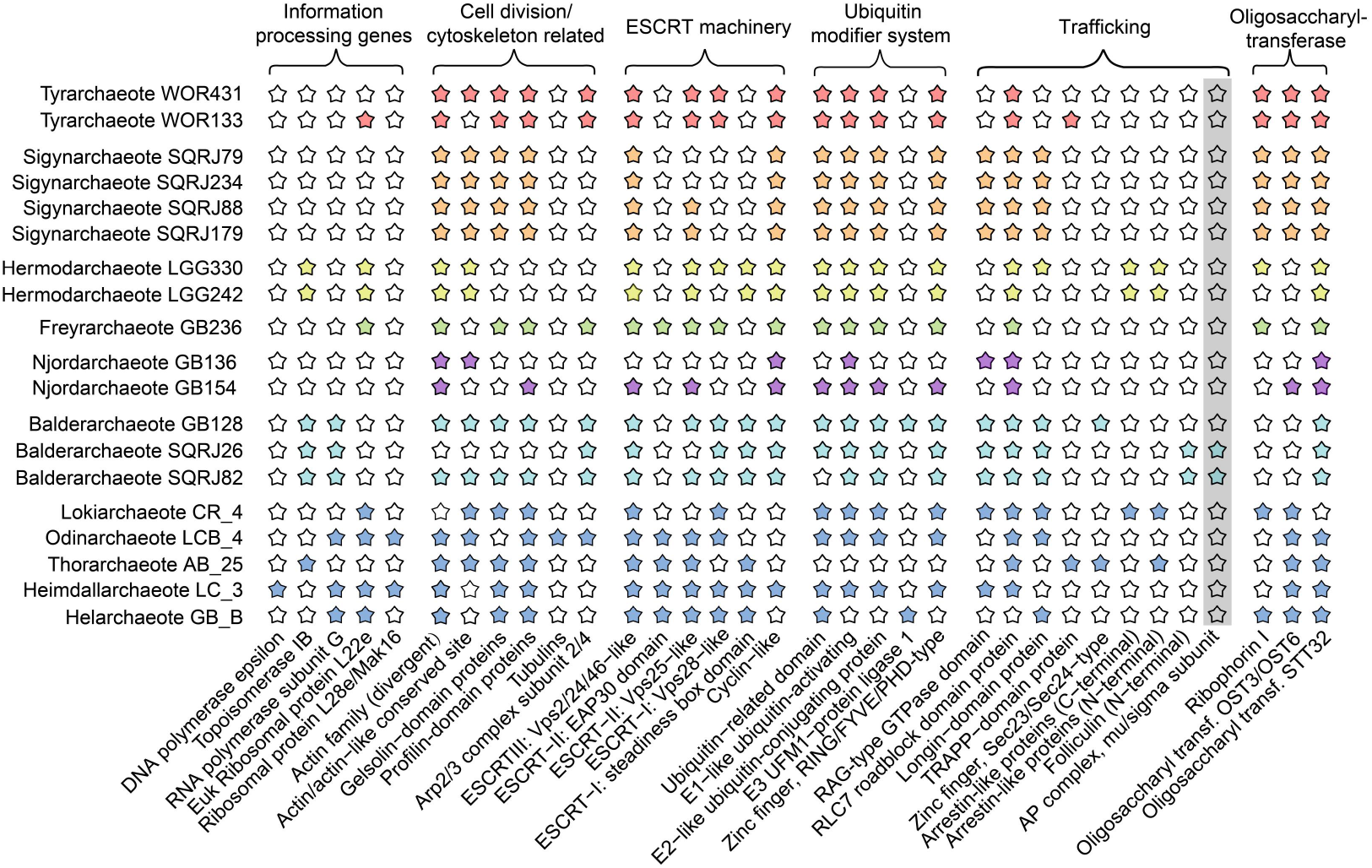
Comparison of the distributions of ESPs in the five new Asgard lineages and other representative Asgard clades. Colored stars indicate the presence of ESPs, whereas empty stars indicate the absence of ESPs. The grey box highlights ESP identified in this study. The new ESP, mu/sigma subunit of AP complex, was detected in Balderarchaeote SQRJ26 and Balderarchaeote SQRJ82.

Ubiquitin-based signaling is an important cellular process in eukaryotes (Raiborg and Stenmark 2009). Previous studies have reported the presence of the related protein domains in Loki-, Odin-, Hel- and Heimdallarchaeota but not in Thorarchaeota (Spang et al., 2015, Seitz et al., 2019, Grau-Bové et al., 2015). Here, we identified ubiquitin system-related protein-encoding genes in nearly all newly assembled MAGs including several ubiquitin-related domains, zinc fingers, ubiquitin-activating enzyme (E1), ubiquitin-conjugating protein (E2), and UFM1-protein ligase 1 (E3), indicating that the ubiquitin system is widespread in Asgard archaea.

The endosomal sorting complex required for transport (ESCRT) machinery consisting of complexes I-III and associated subunits (Spang et al., 2015, Leung et al., 2008, Field and Dacks 2009) were identified in the newly recovered MAGs. Genes coding for Vps28 domain-containing proteins previously found in Loki-, Odin-, Hel- and Heimdallarchaeota (Zaremba-Niedzwiedzka et al., 2017b, Seitz et al., 2019) were also identified in Tyr-, Freyr-, Balder- and Hermodarchaeota but were absent in Sigynarchaeota and Njordarchaeota. The Sigynarchaeota MAGs also lack genes for both EAP30 domain- and steadiness box domain-containing proteins. Notably, all new Asgard MAGs contain cyclin-like protein-encoding genes, whereas ESCRT complexes I-III were only identified in Freyrarchaeota.

ESPs with intracellular trafficking and secretion functions were also identified in MAGs here. However, only Tyrarchaeota contains genes coding for proteins with homology to TRAPP-domain protein and Sec23/24-type protein-encoding gene was only found in Balderarchaeota. All MAGs reported in present study possess genes coding RLC7 roadblock domain protein. Genes coding for both the N- and C-termini of arrestins were found in Hermodarchaeota; the organization resembles one previously reported in the genome of Lokiarchaeote CR-4, in which the C- and N-terminal domain proteins are separated from each other by one gene (Zaremba-Niedzwiedzka et al., 2017b).

We also analyzed the oligosaccharyltransferase (OST) complex in the reconstructed MAGs, and the results showed that OST complex-encoding genes were present in the MAGs of all five Asgard clades. Ribophorin I homolog-encoding genes were found in the Hermod-, Freyr-, Sigyn- and Tyrarchaeota MAGs. Homologs of OST3/6, which have been demonstrated to influence yeast glycosylation efficiency (Schulz et al., 2009), were also identified in several Asgard MAGs, while STT32 subunit protein-encoding genes were detected all MAGs, consistent with previous reports on Loki-, Odin-, Thor-, Hel- and Heimdallarchaeota (Zaremba-Niedzwiedzka et al., 2017b).

In addition to reported ESPs, we identified a potential ESP belonging to mu/sigma subunit of AP (adaptor protein) complex-encoding genes in Balder- and Freyrarchaeota MAGs (Fig. 2). Homologues of mu/sigma subunit of AP complex contain IPR022775 domain. AP complexes are classified into AP-1, AP-2, AP-3, AP-4 and AP-5 and all AP complexes are heterotetramers consisting of two large subunits (adaptins), one medium-sized subunit (mu) and one small-sized subunit (sigma) (Park and Guo 2014, Hirst et al., 2011). AP complexes play a vital role in mediating intracellular membrane trafficking (Tan and Gleeson 2019). Taken together, the identification of potential ESP provides further insight into the origins of eukaryotic cellular complexity.

### Metabolic reconstructions of the five new Asgard lineages

Balderarchaeota MAGs contain genes coding for complete glycolysis via Embden-Meyerhof-Parnas (EMP) pathway, major steps of the tricarboxylic acid (TCA) cycle and β-oxidation pathway, suggesting that members of Balderarchaeota may have potential to metabolize organic compounds including carbohydrates and fatty acids (Fig. 3). The ADP-dependent acetyl-CoA synthetase (ACD) for acetogenesis, which is widely found in archaea (Lazar et al., 2016), was identified in Balderarchaeota MAGs. Meanwhile, phosphate acetyltransferase (Pta) and acetate kinase (Ack) were also found in all Balderarchaeota MAGs (Fig. 3, Supplementary Table 5). Although the *pta* gene was found in Sigynarchaeota as well, all of their MAGs lack the *ack* gene. The Pta/Ack pathway for acetate production, which is common in bacteria, was so far only found in Bathyarchaeota and the methanogenic genus Methanosarcina in archaea (He et al., 2016, Rother and Metcalf 2004) and it is the first case that genes coding for Pta and Ack were discovered in the Asgard archaea. The archaeal *pta/ack* genes were considered HGT from bacteria donors. For example, the genes in Methanosarcina were postulated to acquire from a cellulolytic Clostridia group (Fournier and Gogarten 2008) whereas the *pta/ack* genes donor of Bathyarchaeota was still unclear, possibly transferring independently from unknown clades of Bacteria (He et al., 2016). For Balderarchaeota, the phylogenetic analysis of the *ack* gene sequences revealed that *ack* genes of Balderarchaeota form a monophylogenetic branch (Supplementary Fig. 3), but the phylogenetic tree of *pta* genes shows that the Balderarchaeota clade are within Bathyarchaeota branch (Supplementary Fig. 4), indicating that *pta* genes in Balderarchaeota and Bathyarchaeota probably evolved from the same bacterial donor. Taken together, the Pta/Ack pathway in Balderarchaeota may have acquired from different bacterial donors by two separate HGT events. Additionally, genes coding for the nitrite reductase (NADH) large subunit (*nir*B) were detected in MAGs of Balderarchaeota, implying a potential nitrite reduction capability.

**Figure 3.**
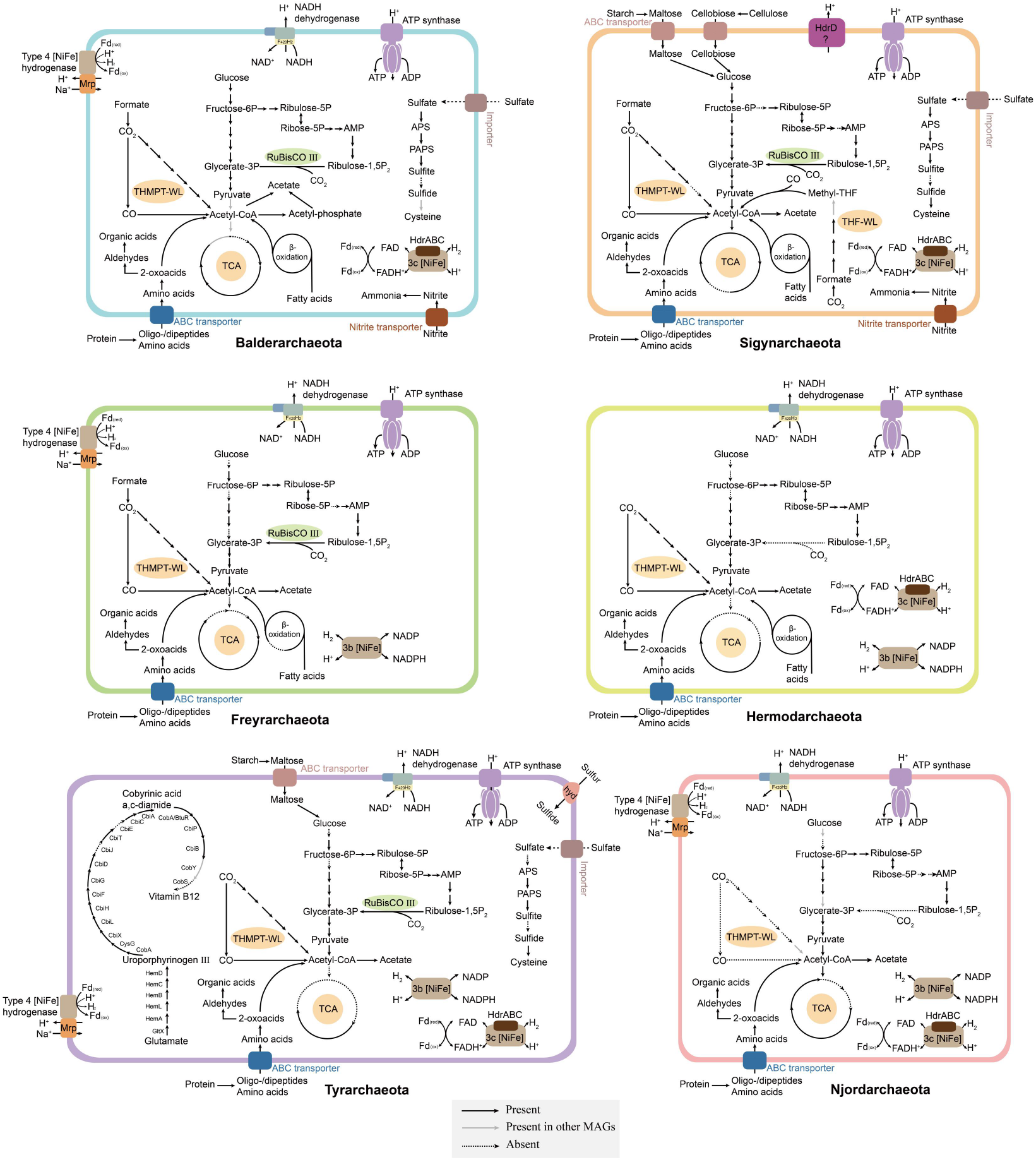
Inferred metabolic pathways of the five new Asgard lineages and the new clade of Hermodarchaeota based on genes identified using the KEGG database and the NCBI NR protein database. A black line indicates that a component/process is present in representative MAGs, a grey line indicates that a component/process is present in other MAGs, and a dashed line indicates that a certain pathway or enzyme is absent from all genomes. The representatives of the different lineages are as follows: Balderarchaeota, SQRJ26; Freyrarchaeota, GB236; Njordarchaeota, GB154; Sigynarchaeota, SQRJ79; and Tyrarchaeota, WOR431; Hermodarchaeota, LGG330. Details about the genes are provided in Supplementary Table 5. Hdr, heterodisulfide reductase; TCA, tricarboxylic acid cycle; THMPT-WL, tetrahydromethanopterin Wood-Ljungdahl pathway; THF-WL, tetrahydrofolate Wood-Ljungdahl; Mrp, Mrp Na^+^/H^+^ antiporters; hyd, sulfhydrogenase; AMP, AMP phosphorylase.

The Wood Ljungdahl (WL) pathway enables organisms to reduce two molecules of CO_2_ to form acetyl-CoA, and then to acetate to produce ATP in anaerobic conditions and Archaea normally utilize tetrahydromethanopterin (THMPT) as C1 carrier. Complete gene sets encoding for the THMPT-WL pathway were found in Freyrarchaeota, suggesting the ability to reduce CO2 through autotrophic acetogenesis (Fig. 3). Tetrahydrofolate (THF) could be as C1 carrier as well, which was generally found in acetogenic bacteria. Sigynarchaeota MAGs contain genes for both types of the WL pathway but 5,10-methylenetetrahydromethanopterin reductase (Mer) which converts 5-methyltetrahydromethanopterin to 5,10-methylenetetrahydromethanopterin was missing in all MAGs identified here, suggesting that Sigynarchaeota probably use the THF-WL pathway for acetate production (Fig. 3, Supplementary Table 5). Sigynarchaeota contain not only all genes responsible for glycolysis but also abundant genes coding for extracellular carbohydrate-degrading enzymes, including α-amylase, cellulase, α-mannosidases and β-glucosidases (Supplementary Table 4), indicating that archaea in Sigynarchaeota have the capacity to degrade complex carbohydrates. Sigynarchaeota MAGs contain neither NADH dehydrogenase nor type 4 [NiFe] hydrogenase (Supplementary Table 5), nevertheless, they probably use membrane-bound heterodisulfide reductase (Hdr) to generate proton motive force as described in their sister-lineage Lokiarchaeota (Spang et al., 2019a).

Among genes associated with glycolysis, the specific enzymes catalyzing these steps is different in the three lineages. In the first step of glycolysis, for example, Sigynarchaeota use ATP-dependent ROK (repressor, open reading frame, kinase) family enzymes while Balderarchaeota utilizes ADP-dependent glucokinase. Likewise, Sigynarchaeota encode ATP-dependent phosphofructokinase (PfkB) but fructose 6-phosphate (F6P) to fructose 1,6-bisphosphate (F1,6P) was catalyzed by ADP-dependent phosphofructokinase (ADP-PFK) in Balderarchaeota.

We also compared differences of metabolic characteristic between the newly identified Hermodarchaeota members and Odinarchaeota since both of them were recovered from high-temperature environments. Hermodarchaeota encodes all enzymes for the complete THMPT-WL pathway, while Odinarchaeota lack several key genes of THMPT-WL pathway. Furthermore, only group 3 [NiFe]-hydrogenases were found in Hermodarchaeota genomes, lacking group 4 [NiFe]- hydrogenases which were identified in Odinarchaeota (Spang et al., 2019a). The presence of THMPT-WL pathway and group 3 [NiFe]-hydrogenases in Hermodarchaeota indicate that they could grow lithoautotrophically by using H_2_ as an electron donor. DnaK-DnaJ-GrpE chaperone system, one of the characteristics of hyperthermophilic archaea (Richter et al., 2010), were found in Hermodarchaeota and Odinarchaeota but genes coding for reverse gyrase were absent in both of them.

In Tyrarchaeota MAGs, glycolysis and TCA pathways are not complete, however, the presence of genes coding for THMPT-WL pathway and group 3 [NiFe] hydrogenases implies its potential to harness energy from H_2_ oxidation, possibly for lithoautotrophic growth, depending on environmental conditions, as suggested for Lokiarchaeota and Thorarchaeota (Spang et al., 2019b, Liu et al., 2018b). Moreover, *de novo* anaerobic cobalamin (vitamin B_12_) biosynthesis pathway was found in Tyrarchaeota (Fig. 3, Supplementary Table 5), suggesting that Tyrarchaeota harbor the potential of cobalamin synthesis. In nature, only limited members of bacteria and archaea possess capacity of *de novo* cobalamin synthesis using one of two alternative pathways: aerobic or anaerobic pathway (Fang et al., 2018). Within Archaea, some members within Euryarchaeota, Thaumarchaeota, Crenarchaeota and Bathyarchaeota have been reported possessing cobalamin synthesizing pathway (Doxey et al., 2015, Pan et al., 2020), whereas only Tyrarchaeota seems to have this ability in the Asgard archaea reported so far.

Njordarchaeota MAGs contain limited genes coding for major carbon metabolic repertoire. Key genes coding for EMP pathway, TCA, the WL pathway and β-oxidation pathway were missing (Fig. 3, Supplementary Table 5). But genes coding for amino acids utilization were found in Njordarchaeota, including aminotransferases and 2-oxoacid:ferredoxin oxidoreductases (the former catalyzes the interconversion of amino acids and 2-oxoacids and the latter oxidates 2-oxoacids to acyl-CoA), indicating that Njordarchaeota has potential to metabolize amino acids. The amino acid carboxylate is transferred to CO_2_ and reducing ferredoxin during 2-oxoacid oxidation, which further could be oxidized into formate and H_2_ by formate dehydrogenase (Fdh) and [NiFe] hydrogenases, respectively (Imachi et al., 2020). [NiFe]-hydrogenases were detected in Njordarchaeota but lacking Fdh, indicating that Njordarchaeota might have a fermentative life style by produce acetate or H_2_ via degrading amino acids. Nevertheless, cultivation experiments are necessary to verify all these predictions described here.

Several models for the origin of eukaryotes have been proposed, based on a metabolic symbiosis between one archaeon and bacterial partner, which was fostered by the discovery of natural syntrophy between *Candidatus* Prometheoarchaeum syntrophicum (that can degrade amino acids to H_2_ or formate) and Deltaproteobacteria (that can utilize H_2_ or Formate and provide amino acids or vitamin B_12_ to partner). Compared with other members of Asgard archaea, Njordarchaeota possess limited pathway for carbon metabolism (Supplementary Table 6), implying that they prone to grow in symbiosis with other organisms to adapt complicated and volatile environment. Combining the phylogenetic affiliation with eukaryotes and metabolic characteristic of Njordarchaeota, we speculate that the archaeal ancestor of eukaryotes probably has potential to degrade amino acids to produce acetate or H_2_ which can further benefit to the bacterial partner, although other additional lifestyles could not be excluded.

## Conclusion

Undoubtedly, the origin of eukaryotes is one of the most important evolutionary events. The discovery of Asgard archaea has boosted the eocyte hypothesis that eukaryotes derive from within archaea because the Asgard archaea possess two remarkable features: robust evolutionary affinity with eukaryotes and various ESPs existence. In the present study, five novel Asgard lineages were discovered based on phylogenetic analyses and AAI value comparison, which significantly expand the phylogenetic and metabolic diversity of the Asgard archaea. Our analyses support a 2-domain tree of life and demonstrate that the eukaryotes lineage cluster with Njordarchaeota alone, suggesting that Njordarchaeota lineage is the closest relatives to eukaryotes than Heimdallarchaeota. Metabolic characteristic of Njordarchaeota shows different carbon metabolic pathways from Heimdallarchaeota that were considered living in anoxic or oxic environment using various organic substrates such as carbohydrates and fatty acids (Spang et al., 2019a, Bulzu et al., 2019, Cai et al., 2020), whereas Njordarchaeota lack both complete glycolysis and WL pathway and the most possibility of metabolism type is using amino acids in anoxic niches. This finding does not contradict to the hypothesis by Spang *et al.* inferring metabolic feature of the archaeal ancestor of eukaryotes (Spang et al., 2019a) that it used organic substrates to produce acetate, formate, H_2_, which might be beneficial for symbiosis. In general, the characterization of additional genomes and continuous efforts to cultivate Asgard archaea will provide additional insights into the evolution of archaea and their potential evolution into eukaryotes. Such insights will enable greater understanding of the ecological and geochemical roles of archaea in Earth’s history.

### Propose type of new taxa

#### *Candidatus* Tyrarchaeum

(Tyr.ar.chae’um. N.L. neut. n. *archaeum* archaeon; N.L. neut. n. *Tyrarchaeum* an archaeon named after Tyr, the god of war in North mythology). Type species: Candidatus Tyrarchaeum oakense.

#### *Candidatus* Tyrarchaeum oakense

(oak’ense N.L. neut. adj. pertaining to white oak river, North Carolina in the United States). This uncultured lineage is represented by the genome “WOR_431” consisting of 2.4 Mbps in 246 contigs with an estimated completeness of 91.56%, an estimated contamination of 2.95% and 20 tRNAs The MAG recovered from white oak river sediment.

#### *Candidatus* Freyrarchaeum

(Freyr.ar.chae’um. N.L. neut. n. *archaeum* archaeon; N.L. neut. n. *Freyrarchaeum* an archaeon named after Freyr, the god of peace in North mythology). Type species: Candidatus Freyrarchaeum guaymasis.

#### *Candidatus* Freyrarchaeum guaymasis

(guayma’sis N.L. neut. adj. pertaining to Guaymas Basin, located in the Gulf of California, México). This uncultured lineage is represented by the genome “GB_236” consisting of 2.0 Mbps in 104 contigs with an estimated completeness of 91.59%, an estimated contamination of 6.54% and 19 tRNAs The MAG recovered from Guaymas Basin sediment.

#### *Candidatus* Sigynarchaeum

(Sigyn.ar.chae’um. N.L. neut. n. *archaeum* archaeon; N.L. neut. n. *Sigynarchaeum* an archaeon named after Sigyn, the god of victory in North mythology). Type species: Candidatus Sigynarchaeum springense.

#### *Candidatus* Sigynarchaeum springense

(spring’ense N.L. neut. adj. pertaining to hot spring, Tengchong, China). This uncultured lineage is represented by the genome “SQRJ_234” consisting of 5.9 Mbps in 269 contigs with an estimated completeness of 91.59%, an estimated contamination of 4.67% and 21 tRNAs The MAG recovered from hot spring sediment.

#### *Candidatus* Balderarchaeum

(Balder.ar.chae’um. N.L. neut. n. *archaeum* archaeon; N.L. neut. n. *Balderarchaeum* an archaeon named after Balder, the god of light in North mythology). Type species: Candidatus Balderarchaeum guaymasis.

#### *Candidatus* Balderarchaeum guaymasis

(guayma’sis N.L. neut. adj. pertaining to Guaymas Basin, located in the Gulf of California, México). This uncultured lineage is represented by the genome “GB_128” consisting of 3.8 Mbps in 131 contigs with an estimated completeness of 97.2%, an estimated contamination of 4.21% and 20 tRNAs The MAG recovered from Guaymas Basin sediment.

#### *Candidatus* Njordarchaeum

(Njord.ar.chae’um. N.L. neut. n. *archaeum* archaeon; N.L. neut. n. *Njordarchaeum* an archaeon named after Njord, the god of seas in North mythology). Type species: Candidatus Njordarchaeum guaymasis.

#### *Candidatus* Njordarchaeum guaymasis

(guayma’sis N.L. neut. adj. pertaining to Guaymas Basin, located in the Gulf of California, México). This uncultured lineage is represented by the genome “GB_154” consisting of 2.1 Mbps in 191 contigs with an estimated completeness of 87.38%, an estimated contamination of 4.67% and 20 tRNAs The MAG recovered from Guaymas Basin sediment.

#### *Candidatus* Tyrarchaeaceae

(Tyr.ar.chae.ace’ae. N.L. neut. n. *Tyrarchaeum*, Candidatus generic name; -aceae ending to denote the family; N.L. fem. pl. n. *Tyrarchaeaceae*, the *Tyrarchaeum* family).

The family is described based on 37 concatenated conserved marker genes phylogeny. The description is the same as that of its sole genus and species. Type genus is *Candidatus* Tyrarchaeum.

#### *Candidatus* Tyrarchaeales

(Tyr.ar.chae.a’les. N.L. neut. n. *Tyrarchaeum*, Candidatus generic name; -ales ending to denote the order; N.L. fem. pl. n. *Tyrarchaeales*, the *Tyrarchaeum* order).

The order is described based on 37 concatenated conserved marker genes phylogeny. The description is the same as that of its sole genus and species. Type genus is *Candidatus* Tyrarchaeum.

#### *Candidatus* Tyrarchaeia

(Tyr.ar.chae’i.a. N.L. neut. n. *Tyrarchaeum*, Candidatus generic name; -ia ending to denote the class; N.L. fem. pl. n. *Tyrarchaeia*, the *Tyrarchaeum* class).

The class is described based on 37 concatenated conserved marker genes phylogeny. The description is the same as that of its sole genus and species. Type genus is *Candidatus* Tyrarchaeum.

#### *Candidatus* Tyrarchaeota

(Tyr.ar.chae.o’ta. N.L. neut. n. *Tyrarchaeum*, Candidatus generic name; -ota ending to denote the phylum; N.L. fem. pl. n. *Tyrarchaeota*, the *Tyrarchaeum* phylum).

The phylum is described based on 37 concatenated conserved marker genes phylogeny. The description is the same as that of its sole genus and species. Type genus is *Candidatus* Tyrarchaeum.

#### *Candidatus* Freyrarchaeaceae

(Freyr.ar.chae.ace’ae. N.L. neut. n. *Freyrarchaeum*, Candidatus generic name; -aceae ending to denote the family; N.L. fem. pl. n. *Freyrarchaeaceae*, the *Freyrarchaeum* family).

The family is described based on 37 concatenated conserved marker genes phylogeny. The description is the same as that of its sole genus and species. Type genus is *Candidatus* Freyrarchaeum.

#### *Candidatus* Freyrarchaeales

(Freyr.ar.chae.a’les. N.L. neut. n. *Freyrarchaeum*, Candidatus generic name; -ales ending to denote the order; N.L. fem. pl. n. *Freyrarchaeales*, the *Freyrarchaeum* order).

The order is described based on 37 concatenated conserved marker genes phylogeny. The description is the same as that of its sole genus and species. Type genus is *Candidatus* Freyrarchaeum.

#### *Candidatus* Freyrarchaeia

(Freyr.ar.chae’i.a. N.L. neut. n. *Freyrarchaeum*, Candidatus generic name; -ia ending to denote the class; N.L. fem. pl. n. *Freyrarchaeia*, the *Freyrarchaeum* class).

The class is described based on 37 concatenated conserved marker genes phylogeny. The description is the same as that of its sole genus and species. Type genus is *Candidatus* Freyrarchaeum.

#### *Candidatus* Freyrarchaeota

(Freyr.ar.chae.o’ta. N.L. neut. n. *Freyrarchaeum*, Candidatus generic name; -ota ending to denote the phylum; N.L. fem. pl. n. *Freyrarchaeota*, the *Freyrarchaeum* phylum).

The phylum is described based on 37 concatenated conserved marker genes phylogeny. The description is the same as that of its sole genus and species. Type genus is *Candidatus* Freyrarchaeum.

#### *Candidatus* Sigynarchaeaceae

(Sigyn.ar.chae.ace’ae. N.L. neut. n. *Sigynarchaeum*, Candidatus generic name; -aceae ending to denote the family; N.L. fem. pl. n. *Sigynarchaeaceae*, the *Sigynarchaeum* family).

The family is described based on 37 concatenated conserved marker genes phylogeny. The description is the same as that of its sole genus and species. Type genus is *Candidatus* Sigynarchaeum.

#### *Candidatus* Sigynarchaeales

(Sigyn.ar.chae.a’les. N.L. neut. n. *Sigynarchaeum*, Candidatus generic name; -ales ending to denote the order; N.L. fem. pl. n. *Sigynarchaeales*, the *Sigynarchaeum* order).

The order is described based on 37 concatenated conserved marker genes phylogeny. The description is the same as that of its sole genus and species. Type genus is *Candidatus* Sigynarchaeum.

#### *Candidatus* Sigynarchaeia

(Sigyn.ar.chae’i.a. N.L. neut. n. *Sigynarchaeum*, Candidatus generic name; -ia ending to denote the class; N.L. fem. pl. n. *Sigynarchaeia*, the *Sigynarchaeum* class).

The class is described based on 37 concatenated conserved marker genes phylogeny. The description is the same as that of its sole genus and species. Type genus is *Candidatus* Sigynarchaeum.

#### *Candidatus* Sigynarchaeota

(Sigyn.ar.chae.o’ta. N.L. neut. n. *Sigynarchaeum*, Candidatus generic name; -ota ending to denote the phylum; N.L. fem. pl. n. *Sigynarchaeota*, the *Sigynarchaeum* phylum).

The phylum is described based on 37 concatenated conserved marker genes phylogeny. The description is the same as that of its sole genus and species. Type genus is *Candidatus* Sigynarchaeum.

#### *Candidatus* Balderarchaeaceae

(Balder.ar.chae.ace’ae. N.L. neut. n. *Balderarchaeum*, Candidatus generic name; -aceae ending to denote the family; N.L. fem. pl. n. *Balderarchaeaceae*, the *Balderarchaeum* family).

The family is described based on 37 concatenated conserved marker genes phylogeny. The description is the same as that of its sole genus and species. Type genus is *Candidatus* Balderarchaeum.

#### *Candidatus* Balderarchaeales

(Balder.ar.chae.a’les. N.L. neut. n. *Balderarchaeum*, Candidatus generic name; -ales ending to denote the order; N.L. fem. pl. n. *Balderarchaeales*, the *Balderarchaeum* order).

The order is described based on 37 concatenated conserved marker genes phylogeny. The description is the same as that of its sole genus and species. Type genus is *Candidatus* Balderarchaeum.

#### *Candidatus* Balderarchaeia

(Balder.ar.chae’i.a. N.L. neut. n. *Balderarchaeum*, Candidatus generic name; -ia ending to denote the class; N.L. fem. pl. n. *Balderarchaeia*, the *Balderarchaeum* class).

The class is described based on 37 concatenated conserved marker genes phylogeny. The description is the same as that of its sole genus and species. Type genus is *Candidatus* Balderarchaeum.

#### *Candidatus* Balderarchaeota

(Balder.ar.chae.o’ta. N.L. neut. n. *Balderarchaeum*, Candidatus generic name; -ota ending to denote the phylum; N.L. fem. pl. n. *Balderarchaeota*, the *Balderarchaeum* phylum).

The phylum is described based on 37 concatenated conserved marker genes phylogeny. The description is the same as that of its sole genus and species. Type genus is *Candidatus* Balderarchaeum.

#### *Candidatus* Njordarchaeaceae

(Njord.ar.chae.ace’ae. N.L. neut. n. *Njordarchaeum*, Candidatus generic name; -aceae ending to denote the family; N.L. fem. pl. n. *Njordarchaeaceae*, the *Njordarchaeum* family).

The family is described based on 37 concatenated conserved marker genes phylogeny. The description is the same as that of its sole genus and species. Type genus is *Candidatus* Njordarchaeum.

#### *Candidatus* Njordarchaeales

(Njord.ar.chae.a’les. N.L. neut. n. *Njordarchaeum*, Candidatus generic name; -ales ending to denote the order; N.L. fem. pl. n. *Njordarchaeales*, the *Njordarchaeum* order).

The order is described based on 37 concatenated conserved marker genes phylogeny. The description is the same as that of its sole genus and species. Type genus is *Candidatus* Njordarchaeum.

#### *Candidatus* Njordarchaeia

(Njord.ar.chae’i.a. N.L. neut. n. *Njordarchaeum*, Candidatus generic name; -ia ending to denote the class; N.L. fem. pl. n. *Njordarchaeia*, the *Njordarchaeum* class).

The class is described based on 37 concatenated conserved marker genes phylogeny. The description is the same as that of its sole genus and species. Type genus is *Candidatus* Njordarchaeum.

#### *Candidatus* Njordarchaeota

(Njord.ar.chae.o’ta. N.L. neut. n. *Njordarchaeum*, Candidatus generic name; -ota ending to denote the phylum; N.L. fem. pl. n. *Njordarchaeota*, the *Njordarchaeum* phylum).

The phylum is described based on 37 concatenated conserved marker genes phylogeny. The description is the same as that of its sole genus and species. Type genus is *Candidatus* Njordarchaeum.

## Methods

### Sampling and processing

Detailed methods for collection, DNA extraction, and metagenome sequencing of Guaymas Basin samples has been described in previous study (Feng et al., 2019). Six sediment samples of hot spring were taken from Tengchong, Yunnan, China on September, 2019 (24.95°N, 98.44°E). DNA was extracted from 10 g of each sample by using PowerSoil DNA Isolation Kit (Mo Bio). Metagenomic sequence data for the six samples were generated using Illumina HiSeq 2500 instruments.

### Data collection

Asgard archaea are distributed mainly in different environmental sediments, including estuary sediments (Seitz et al., 2016), mangrove sediments (Liu et al., 2018a), hydrothermal sediments (Dombrowski et al., 2018), hot spring sediments (Zaremba-Niedzwiedzka et al., 2017a), marine sediments (Spang et al., 2015), and freshwater sediments (Narrowe et al., 2018). According to the environmental distributions of Asgard archaea, metagenomic data were collected and downloaded from the SRA database (https://www.ncbi.nlm.nih.gov/sra/).

### Metagenomic assembly and genomic binning

The raw reads were trimmed using Trimmomatic (v.0.38) (Bolger et al., 2014) to remove adapters and low-quality reads. After trimming, the reads of each sample were *de novo* assembled using Megahit (v.1.2.5) (Li et al., 2015) with a k-step of 6. Samples from the same location or similar environments were assembled together. Contigs were binned separately using MetaBAT (v.2.12.1) (Kang et al., 2019), MaxBin (v.2.2.7) (Wu et al., 2016), and Concoct (v.1.1.0) (Alneberg et al., 2014) with the default parameters, and the initial taxonomic classification of each MAGs was performed using GTDB-Tk (v.1.2.0) (Chaumeil et al., 2019) to extract Asgard MAGs. The completeness and contamination of Asgard MAGs were evaluated with the CheckM lineage_wf workflow (v.1.0.12) (Parks et al., 2015). Finally, Asgard MAGs with completeness above 50% and contamination below 10% were selected for further analyses, and Prodigal (v.2.6.1) (Hyatt et al., 2010) was used to predict protein-coding genes for these selected Asgard MAGs.

### Phylogenetic analyses of Asgard MAGs

To determine the exact phylogenetic affiliations in the Asgard superphylum, 37 conserved marker genes were selected as described in the literature (Wang et al., 2019, Jay et al., 2018). Homologs of the 37 conserved marker proteins were identified using Diamond (v.2.0.4) (Buchfink et al., 2015). Each dataset of marker proteins was aligned with MAFFT-L-INS-i (v.7.313) (Katoh and Standley 2013) and trimmed by trimAl (v.1.4.22) (Capella-Gutiérrez et al., 2009) with the “automated1” option. Maximum-likelihood phylogenies for the 37 conserved marker proteins was built using IQ-TREE (v.2.0.5) (Nguyen et al., 2015) under the model “LG+F+G4+C60”. The support values were calculated using 1000 ultrafast bootstraps.

### Phylogenetic tree of life

To confirm the phylogenetic affiliations of eukaryotes and the novel Asgard lineages, 21 taxonomic marker genes shared among three domains selected by Williams (Williams et al., 2020) were used for phylogenetic analyses. Single-gene trees were inferred for all the markers using IQ-TREE with the LG+G4+F model to exclude phylogenetic artefacts such as long branch attraction, HGT (eukaryotic genes falling into bacterial clade or scattering in archaeal clade were considered as HGT) and a BLASTp inspection was further performed to identify all eukaryotic genes that originated from the nuclear genome and to remove genes of mitochondrial or chloroplastic origin. The maximum-likelihood tree was built with IQ-TREE under the LG+C60+F+G4 model with 1000 ultrafast bootstraps.

### Identification of ESPs

All predicted proteins encoded by the MAGs of the five novel Asgard lineages and Hermodarchaeota were analyzed using InterProScan (Jones et al., 2014) (v.5.47-82.0) with default parameters to annotate protein domains and were assigned to archaeal clusters of orthologous genes (arCOGs) (Makarova et al., 2015) by eggnog-mapper (v. 2.0.1b) (Huerta-Cepas et al., 2019) with default settings.. The lists of InterPro accession numbers (IPRs) and arCOG identifiers previously published by Zaremba-Niedzwiedzka *et al.* (Zaremba-Niedzwiedzka et al., 2017a) and Bulzu *et al.* (Bulzu et al., 2019) were used to identify potential ESPs. Some key words related to eukaryote-specific processes or cell structures were used to search annotation information of interProScan to identify potential ESPs previously not reported in Asgard archaea. Several candidate ESPs were further examined using HHpred (Zimmermann et al., 2018) with default parameters.

### Metabolic reconstruction

The proteome of each MAG reported in the present study was uploaded to the KEGG Automatic Annotation Server (KAAS) (Moriya et al., 2007) and run with several settings: the GHOSTX, Prokaryotes and Bidirectional Best Hit (BBH) settings. Additionally, proteins were queried against the nonredundant (NR) protein database (downloaded from NCBI on February 2020) using the Diamond (v.2.0.4) BLASTp search (e-value cutoff <1e−5). Metabolic pathways were reconstructed based on combination of the NR annotations, protein domain information and KEGG Ontology (KO) numbers.

The dbCAN2 (Zhang et al., 2018) web server was used to identify carbohydrate-degrading enzymes with the default settings. The putative large subunits of [NiFe] hydrogenases were identified by querying against a local database based on HydDB (Søndergaard et al., 2016) using Diamond (v.2.0.4) with an E-value cutoff of 1e^−20^ and sequences containing CxxC motifs in both N-terminal and C-terminal were considered as hydrogenases. Additionally, a local MEROPS database (downloaded September 2020) (Rawlings et al., 2016) searched for peptidases by Diamond (v.2.0.4) with an E-value cutoff of 1×10^−20^, and PSORT (v.3.0.2) was used to identify protein localization (Horton et al., 2007).

### Calculation of ANI and average AAI

The ANI and AAI values were calculated using OrthoANI (v.1.2) (Lee et al., 2016) and CompareM (https://github.com/dparks1134/CompareM), respectively, with the default parameters.

## Acknowledgments

We are grateful for Dr. Tom A. William for his suggestions regarding phylogenetic analysis. We thank Brett Baker for allowing us use their metagenomic data freely. These sequence data were produced by the US Department of Energy Joint Genome Institute http://www.jgi.doe.gov/ in collaboration with the user community.

## Data availability

The genomes of Asgard archaea generated in this study have been made available at the eLibrary of Microbial Systematics and Genomics (eLMSG; https://www.biosino.org/elmsg/index) under accession numbers LMSG_G000000610.1 - LMSG_G000000611.1 for Njordarchaeota, LMSG_G000000612.1 - LMSG_G000000613.1 for Tyrarchaeota, LMSG_G000000614.1 - LMSG_G000000616.1 and LMSG_G000000628.1 for Sigynarchaeota, LMSG_G000000617.1 - LMSG_G000000618.1 for Hermodarchaeota, LMSG_G000000622.1 for Freyrarchaeota, LMSG_G000000625.1 - LMSG_G000000627.1 for Sigynarchaeota.

The initial phylogenetic trees have been deposited at figshare and can be accessed at the following link: https://figshare.com/s/c20d9eccb7e4591b429c.

## Funding

This work was supported by the Natural Science Foundation of China (Grant No. 91751205, 41525011), the National Key Research and Development Project of China (Grant No. 2016YFA0601102), the Senior User Project of RV KEXUE (KEXUE2019GZ06).

## Author contributions

R.Z.X., Y.Z.W. and F.P.W. conceived the study. R.Z.X., Y.Z.W., D.Y.H., H.J.L., H.N.H., L.Y.L. and X.X.Z. analyzed the data. R.Z.X., Y.Z.W. and F.P.W. wrote the paper.

## Compliance and ethics

The author(s) declare that they have no conflicts of interest.

## Notes

### Competing Interest Statement

The authors have declared no competing interest.

## References

Embley T M, Martin W. 2006. Eukaryotic evolution, changes and challenges. Nature, 440, 623–630.

López-García P, Moreira D. 2015. Open questions on the origin of eukaryotes. Trends in ecology & evolution, 30, 697–708.

Rochette N C, Brochier-Armanet C, Gouy M. 2014. Phylogenomic test of the hypotheses for the evolutionary origin of eukaryotes. Molecular biology and evolution, 31, 832–845.

Lopez-Garcia P, Moreira D. 2006. Selective forces for the origin of the eukaryotic nucleus. Bioessays, 28, 525–533. doi,10.1002/bies.20413.

López-Garcia P, Moreira D. 1999. Metabolic symbiosis at the origin of eukaryotes. Trends in biochemical sciences, 24, 88–93.

Martin W F, Garg S, Zimorski V. 2015. Endosymbiotic theories for eukaryote origin. Philosophical Transactions of the Royal Society B: Biological Sciences, 370, 20140330.

Esser C, Ahmadinejad N, Wiegand C, Rotte C, Sebastiani F, Gelius-Dietrich G, Henze K, Kretschmann E, Richly E, Leister D. 2004. A genome phylogeny for mitochondria among α-proteobacteria and a predominantly eubacterial ancestry of yeast nuclear genes. Molecular biology and evolution, 21, 1643–1660.

Moreira D, López-García P. 1998. Symbiosis between methanogenic archaea and δ-proteobacteria as the origin of eukaryotes: the syntrophic hypothesis. Journal of Molecular Evolution, 47, 517–530.

Spang A, Saw J H, Jorgensen S L, Zaremba-Niedzwiedzka K, Martijn J, Lind A E, van Eijk R, Schleper C, Guy L, Ettema T J G. 2015. Complex archaea that bridge the gap between prokaryotes and eukaryotes. Nature, 521, 173–179. doi,10.1038/nature14447.

Zaremba-Niedzwiedzka K, Caceres E F, Saw J H, Bäckström D, Juzokaite L, Vancaester E, Seitz K W, Anantharaman K, Starnawski P, Kjeldsen K U. 2017a. Asgard archaea illuminate the origin of eukaryotic cellular complexity. Nature, 541, 353.

Spang A, Stairs C W, Dombrowski N, Eme L, Lombard J, Caceres E F, Greening C, Baker B J, Ettema T J. 2019a. Proposal of the reverse flow model for the origin of the eukaryotic cell based on comparative analyses of Asgard archaeal metabolism. Nature microbiology, 1.

Williams T A, Foster P G, Cox C J, Embley T M. 2013. An archaeal origin of eukaryotes supports only two primary domains of life. Nature, 504, 231–236.

Woese C R, Kandler O, Wheelis M L. 1990. Towards a natural system of organisms: proposal for the domains Archaea, Bacteria, and Eucarya. Proceedings of the National Academy of Sciences, 87, 4576–4579.

Williams T A, Cox C J, Foster P G, Szollosi G J, Embley T M. 2020. Phylogenomics provides robust support for a two-domains tree of life. Nat Ecol Evol, 4, 138–147. doi,10.1038/s41559-019-1040-x.

Hartman H, Fedorov A. 2002. The origin of the eukaryotic cell: a genomic investigation. Proceedings of the National Academy of Sciences, 99, 1420–1425.

Bulzu P-A, Andrei A-Ş, Salcher M M, Mehrshad M, Inoue K, Kandori H, Beja O, Ghai R, Banciu H L. 2019. Casting light on Asgardarchaeota metabolism in a sunlit microoxic niche. Nature microbiology, 4, 1129–1137.

Liu Y, Zhou Z, Pan J, Baker B J, Gu J-D, Li M. 2018a. Comparative genomic inference suggests mixotrophic lifestyle for Thorarchaeota. The ISME journal, 12, 1021–1031.

Seitz K W, Lazar C S, Hinrichs K-U, Teske A P, Baker B J. 2016. Genomic reconstruction of a novel, deeply branched sediment archaeal phylum with pathways for acetogenesis and sulfur reduction. The ISME journal, 10, 1696–1705.

Wong H L, White R A, Visscher P T, Charlesworth J C, Vázquez-Campos X, Burns B P. 2018. Disentangling the drivers of functional complexity at the metagenomic level in Shark Bay microbial mat microbiomes. The ISME journal, 12, 2619–2639.

MacLeod F, Kindler G S, Wong H L, Chen R, Burns B P. 2019. Asgard archaea: diversity, function, and evolutionary implications in a range of microbiomes. AIMS microbiology, 5, 48.

Imachi H, Nobu M K, Nakahara N, Morono Y, Ogawara M, Takaki Y, Takano Y, Uematsu K, Ikuta T, Ito M, Matsui Y, Miyazaki M, Murata K, Saito Y, Sakai S, Song C, Tasumi E, Yamanaka Y, Yamaguchi T, Kamagata Y, Tamaki H, Takai K. 2020. Isolation of an archaeon at the prokaryote-eukaryote interface. Nature, 577, 519–525. doi,10.1038/s41586-019-1916-6.

Akil C, Tran L T, Orhant-Prioux M, Baskaran Y, Manser E, Blanchoin L, Robinson R C. 2020. Insights into the evolution of regulated actin dynamics via characterization of primitive gelsolin/cofilin proteins from Asgard archaea. Proceedings of the National Academy of Sciences.

Zhang R-Y, Zou B, Yan Y-W, Jeon C O, Li M, Cai M, Quan Z-X. 2020. Design of targeted primers based on 16S rRNA sequences in meta-transcriptomic datasets and identification of a novel taxonomic group in the Asgard archaea. BMC microbiology, 20, 25.

Wang Y, Wegener G, Hou J, Wang F, Xiao X. 2019. Expanding anaerobic alkane metabolism in the domain of Archaea. Nat Microbiol, 4, 595–602. doi,10.1038/s41564-019-0364-2.

Zaremba-Niedzwiedzka K, Caceres E F, Saw J H, Backstrom D, Juzokaite L, Vancaester E, Seitz K W, Anantharaman K, Starnawski P, Kjeldsen K U, Stott M B, Nunoura T, Banfield J F, Schramm A, Baker B J, Spang A, Ettema T J. 2017b. Asgard archaea illuminate the origin of eukaryotic cellular complexity. Nature, 541, 353–358. doi,10.1038/nature21031.

Zhang J W, Dong H P, Hou L J, Liu Y, Ou Y F, Zheng Y L, Han P, Liang X, Yin G Y, Wu D M, Liu M, Li M., 2021. Newly discovered Asgard archaea Hermodarchaeota potentially degrade alkanes and aromatics via alkyl/benzyl-succinate synthase and benzoyl-CoA pathway. ISME J. doi,10.1038/s41396-020-00890-x.

Luo C, Rodriguez-r L M, Konstantinidis K T. 2014. MyTaxa: an advanced taxonomic classifier for genomic and metagenomic sequences. Nucleic acids research, 42, e73–e73.

Seitz K W, Dombrowski N, Eme L, Spang A, Lombard J, Sieber J R, Teske A P, Ettema T J G, Baker B J. 2019. Asgard archaea capable of anaerobic hydrocarbon cycling. Nat Commun, 10, 1822. doi,10.1038/s41467-019-09364-x.

Raiborg C, Stenmark H. 2009. The ESCRT machinery in endosomal sorting of ubiquitylated membrane proteins. Nature, 458, 445–452.

Grau-Bové X, Sebé-Pedrós A, Ruiz-Trillo I. 2015. The Eukaryotic Ancestor Had a Complex Ubiquitin Signaling System of Archaeal Origin. Molecular Biology and Evolution, 32, 726–739. doi,10.1093/molbev/msu334.

Leung K F, Dacks J B, Field M C. 2008. Evolution of the multivesicular body ESCRT machinery; retention across the eukaryotic lineage. Traffic, 9, 1698–1716. doi,10.1111/j.1600-0854.2008.00797.x.

Field M C, Dacks J B. 2009. First and last ancestors: reconstructing evolution of the endomembrane system with ESCRTs, vesicle coat proteins, and nuclear pore complexes. Curr Opin Cell Biol, 21, 4–13. doi,10.1016/j.ceb.2008.12.004.

Schulz B L, Stirnimann C U, Grimshaw J P, Brozzo M S, Fritsch F, Mohorko E, Capitani G, Glockshuber R, Grutter M G, Aebi M. 2009. Oxidoreductase activity of oligosaccharyltransferase subunits Ost3p and Ost6p defines site-specific glycosylation efficiency. Proc Natl Acad Sci U S A, 106, 11061–11066. doi,10.1073/pnas.0812515106.

Park S Y, Guo X. 2014. Adaptor protein complexes and intracellular transport. Bioscience reports, 34.

Hirst J, Barlow L D, Francisco G C, Sahlender D A, Seaman M N, Dacks J B, Robinson M S. 2011. The fifth adaptor protein complex. PLoS Biol, 9, e1001170.

Tan J Z A, Gleeson P A. 2019. Cargo sorting at the trans-Golgi network for shunting into specific transport routes: role of Arf small G proteins and adaptor complexes. Cells, 8, 531.

Lazar C S, Baker B J, Seitz K, Hyde A S, Dick G J, Hinrichs K U, Teske A P. 2016. Genomic evidence for distinct carbon substrate preferences and ecological niches of B athyarchaeota in estuarine sediments. Environmental microbiology, 18, 1200–1211.

He Y, Li M, Perumal V, Feng X, Fang J, Xie J, Sievert S, Wang F. 2016. Genomic and enzymatic evidence for acetogenesis among multiple lineages of the archaeal phylum Bathyarchaeota widespread in marine sediments. Nature microbiology, 1, 1–9.

Rother M, Metcalf W W. 2004. Anaerobic growth of Methanosarcina acetivorans C2A on carbon monoxide: an unusual way of life for a methanogenic archaeon. Proceedings of the National Academy of Sciences, 101, 16929–16934.

Fournier G P, Gogarten J P. 2008. Evolution of acetoclastic methanogenesis in Methanosarcina via horizontal gene transfer from cellulolytic Clostridia. Journal of bacteriology, 190, 1124–1127.

Sousa F L, Martin W F. 2014. Biochemical fossils of the ancient transition from geoenergetics to bioenergetics in prokaryotic one carbon compound metabolism. Biochimica et Biophysica Acta (BBA)-Bioenergetics, 1837, 964–981.

Ma K, Schicho R N, Kelly R M, Adams M W. 1993. Hydrogenase of the hyperthermophile Pyrococcus furiosus is an elemental sulfur reductase or sulfhydrogenase: evidence for a sulfur-reducing hydrogenase ancestor. Proc Natl Acad Sci U S A, 90, 5341–5344. doi,10.1073/pnas.90.11.5341.

Richter K, Haslbeck M, Buchner J., 2010. The heat shock response: life on the verge of death. Molecular Cell, 40, 253–266.

Spang A, Stairs C W, Dombrowski N, Eme L, Lombard J, Caceres E F, Greening C, Baker B J, Ettema T J G. 2019b. Proposal of the reverse flow model for the origin of the eukaryotic cell based on comparative analyses of Asgard archaeal metabolism. Nat Microbiol, 4, 1138–1148. doi,10.1038/s41564-019-0406-9.

Liu Y, Zhou Z, Pan J, Baker B J, Gu J D, Li M. 2018b. Comparative genomic inference suggests mixotrophic lifestyle for Thorarchaeota. ISME J, 12, 1021–1031. doi,10.1038/s41396-018-0060-x.

Fang H, Li D, Kang J, Jiang P, Sun J, Zhang D., 2018. Metabolic engineering of Escherichia coli for de novo biosynthesis of vitamin B 12. Nature communications, 9, 1–12.

Doxey A C, Kurtz D A, Lynch M D, Sauder L A, Neufeld J D. 2015. Aquatic metagenomes implicate Thaumarchaeota in global cobalamin production. The ISME journal, 9, 461–471.

Pan J, Zhou Z, Béjà O, Cai M, Yang Y, Liu Y, Gu J-D, Li M., 2020. Genomic and transcriptomic evidence of light-sensing, porphyrin biosynthesis, Calvin-Benson-Bassham cycle, and urea production in Bathyarchaeota. Microbiome, 8, 1–12.

Cai M, Liu Y, Yin X, Zhou Z, Friedrich M W, Richter-Heitmann T, Nimzyk R, Kulkarni A, Wang X, Li W., 2020. Diverse Asgard archaea including the novel phylum Gerdarchaeota participate in organic matter degradation. Science China Life Sciences, 1–12.

Feng X, Wang Y, Zubin R, Wang F., 2019. Core metabolic features and hot origin of Bathyarchaeota. Engineering, 5, 498–504.

Dombrowski N, Teske A P, Baker B J. 2018. Expansive microbial metabolic versatility and biodiversity in dynamic Guaymas Basin hydrothermal sediments. Nature communications, 9, 1–13.

Narrowe A B, Spang A, Stairs C W, Caceres E F, Baker B J, Miller C S, Ettema T J. 2018. Complex evolutionary history of translation Elongation Factor 2 and diphthamide biosynthesis in Archaea and parabasalids. Genome biology and evolution, 10, 2380–2393.

Bolger A M, Lohse M, Usadel B. 2014. Trimmomatic: a flexible trimmer for Illumina sequence data. Bioinformatics, 30, 2114–2120.

Li D, Liu C-M, Luo R, Sadakane K, Lam T-W. 2015. MEGAHIT: an ultra-fast single-node solution for large and complex metagenomics assembly via succinct de Bruijn graph. Bioinformatics, 31, 1674–1676.

Kang D D, Li F, Kirton E, Thomas A, Egan R, An H, Wang Z. 2019. MetaBAT 2: an adaptive binning algorithm for robust and efficient genome reconstruction from metagenome assemblies. PeerJ, 7, e7359.

Wu Y-W, Simmons B A, Singer S W. 2016. MaxBin 2.0: an automated binning algorithm to recover genomes from multiple metagenomic datasets. Bioinformatics, 32, 605–607.

Alneberg J, Bjarnason B S, De Bruijn I, Schirmer M, Quick J, Ijaz U Z, Lahti L, Loman N J, Andersson A F, Quince C., 2014. Binning metagenomic contigs by coverage and composition. Nature methods, 11, 1144–1146.

Chaumeil P A, Mussig A J, Hugenholtz P, Parks D H. 2019. GTDB-Tk: a toolkit to classify genomes with the Genome Taxonomy Database. Bioinformatics. doi,10.1093/bioinformatics/btz848.

Parks D H, Imelfort M, Skennerton C T, Hugenholtz P, Tyson G W. 2015. CheckM: assessing the quality of microbial genomes recovered from isolates, single cells, and metagenomes. Genome research, 25, 1043–1055.

Hyatt D, Chen G-L, LoCascio P F, Land M L, Larimer F W, Hauser L J. 2010. Prodigal: prokaryotic gene recognition and translation initiation site identification. BMC bioinformatics, 11, 119.

Jay Z J, Beam J P, Dlakić M, Rusch D B, Kozubal M A, Inskeep W P. 2018. Marsarchaeota are an aerobic archaeal lineage abundant in geothermal iron oxide microbial mats. Nature microbiology, 3, 732–740.

Buchfink B, Xie C, Huson D H. 2015. Fast and sensitive protein alignment using DIAMOND. Nature methods, 12, 59–60.

Katoh K, Standley D M. 2013. MAFFT multiple sequence alignment software version 7: improvements in performance and usability. Molecular biology and evolution, 30, 772–780.

Capella-Gutiérrez S, Silla-Martínez J M, Gabaldón T. 2009. trimAl: a tool for automated alignment trimming in large-scale phylogenetic analyses. Bioinformatics, 25, 1972–1973.

Nguyen L-T, Schmidt H A, Von Haeseler A, Minh B Q. 2015. IQ-TREE: a fast and effective stochastic algorithm for estimating maximum-likelihood phylogenies. Molecular biology and evolution, 32, 268–274.

Jones P, Binns D, Chang H-Y, Fraser M, Li W, McAnulla C, McWilliam H, Maslen J, Mitchell A, Nuka G., 2014. InterProScan 5: genome-scale protein function classification. Bioinformatics, 30, 1236–1240.

Makarova K S, Wolf Y I, Koonin E V. 2015. Archaeal clusters of orthologous genes (arCOGs): an update and application for analysis of shared features between Thermococcales, Methanococcales, and Methanobacteriales. Life, 5, 818–840.

Huerta-Cepas J, Szklarczyk D, Heller D, Hernández-Plaza A, Forslund S K, Cook H, Mende D R, Letunic I, Rattei T, Jensen L J. 2019. eggNOG 5.0: a hierarchical, functionally and phylogenetically annotated orthology resource based on 5090 organisms and 2502 viruses. Nucleic acids research, 47, D309–D314.

Zimmermann L, Stephens A, Nam S-Z, Rau D, Kübler J, Lozajic M, Gabler F, Söding J, Lupas A N, Alva V. 2018. A completely reimplemented MPI bioinformatics toolkit with a new HHpred server at its core. Journal of molecular biology, 430, 2237–2243.

Moriya Y, Itoh M, Okuda S, Yoshizawa A C, Kanehisa M. 2007. KAAS: an automatic genome annotation and pathway reconstruction server. Nucleic acids research, 35, W182–W185.

Zhang H, Yohe T, Huang L, Entwistle S, Wu P, Yang Z, Busk P K, Xu Y, Yin Y., 2018. dbCAN2: a meta server for automated carbohydrate-active enzyme annotation. Nucleic acids research, 46, W95–W101.

Søndergaard D, Pedersen C N, Greening C. 2016. HydDB: a web tool for hydrogenase classification and analysis. Scientific reports, 6, 1–8.

Rawlings N D, Barrett A J, Finn R. 2016. Twenty years of the MEROPS database of proteolytic enzymes, their substrates and inhibitors. Nucleic acids research, 44, D343–D350.

Horton P, Park K-J, Obayashi T, Fujita N, Harada H, Adams-Collier C, Nakai K. 2007. WoLF PSORT: protein localization predictor. Nucleic acids research, 35, W585–W587.

Lee I, Kim Y O, Park S-C, Chun J. 2016. OrthoANI: an improved algorithm and software for calculating average nucleotide identity. International journal of systematic and evolutionary microbiology, 66, 1100–1103.

