## Supplemental materials for "Expanding Asgard members in the domain of Archaea shed new light on the origin of eukaryotes" (from another version of this preprint: version 1, 2021-04-02)

### Supplementary Figures

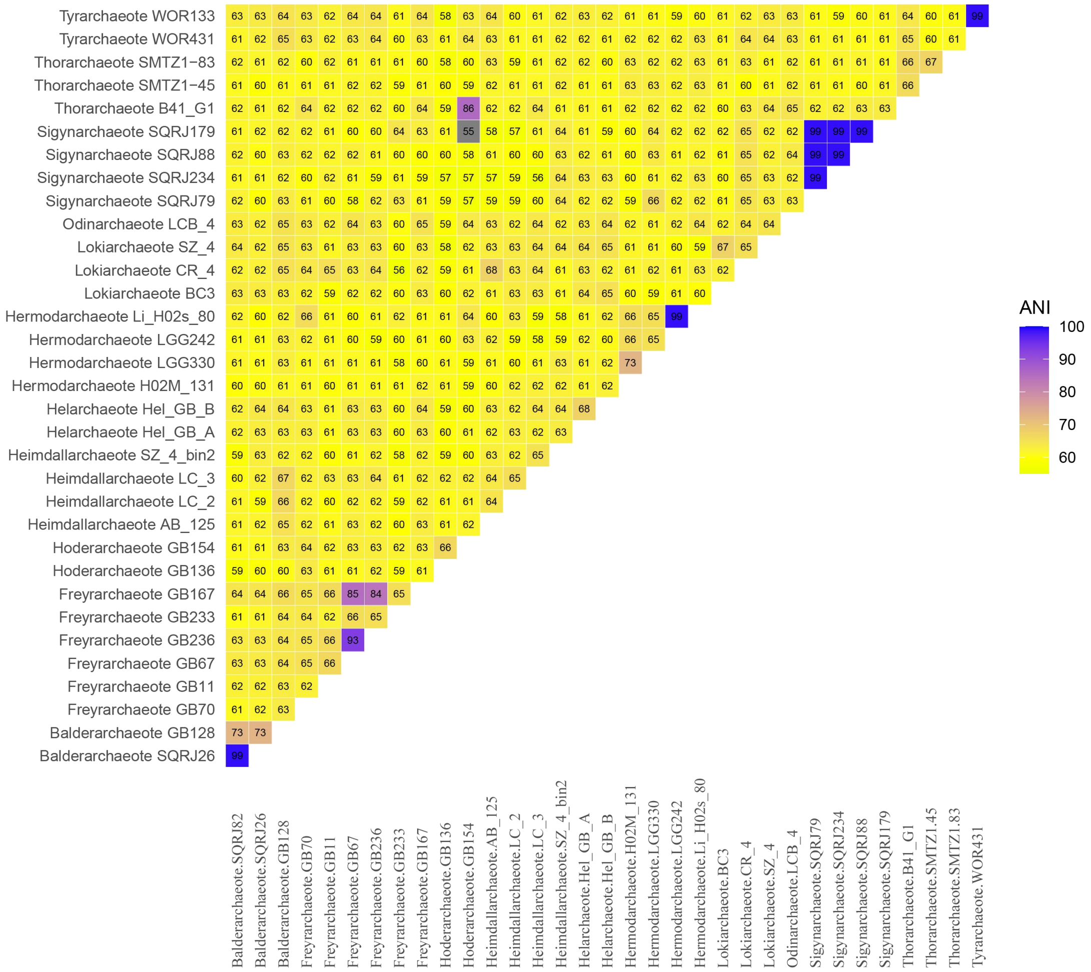


**Supplementary Figure 1.** Average nucleotide identity (ANI) values of the six novel Asgard archaea with the known Asgard archaea as calculated by OrthoANI.

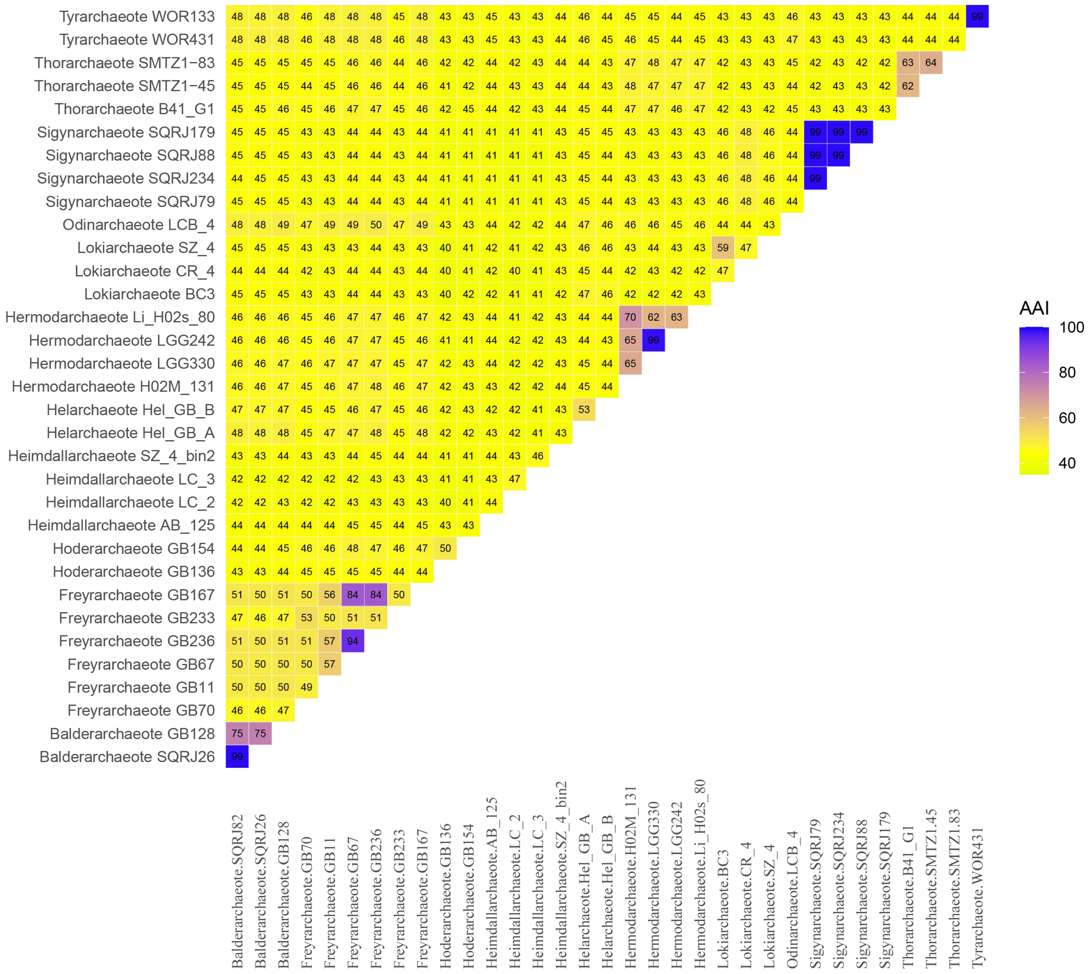

**Supplementary Figure 2.** Amino acid identity (AAI) values of the six novel Asgard archaea with the known Asgard archaea as calculated by CompareM.

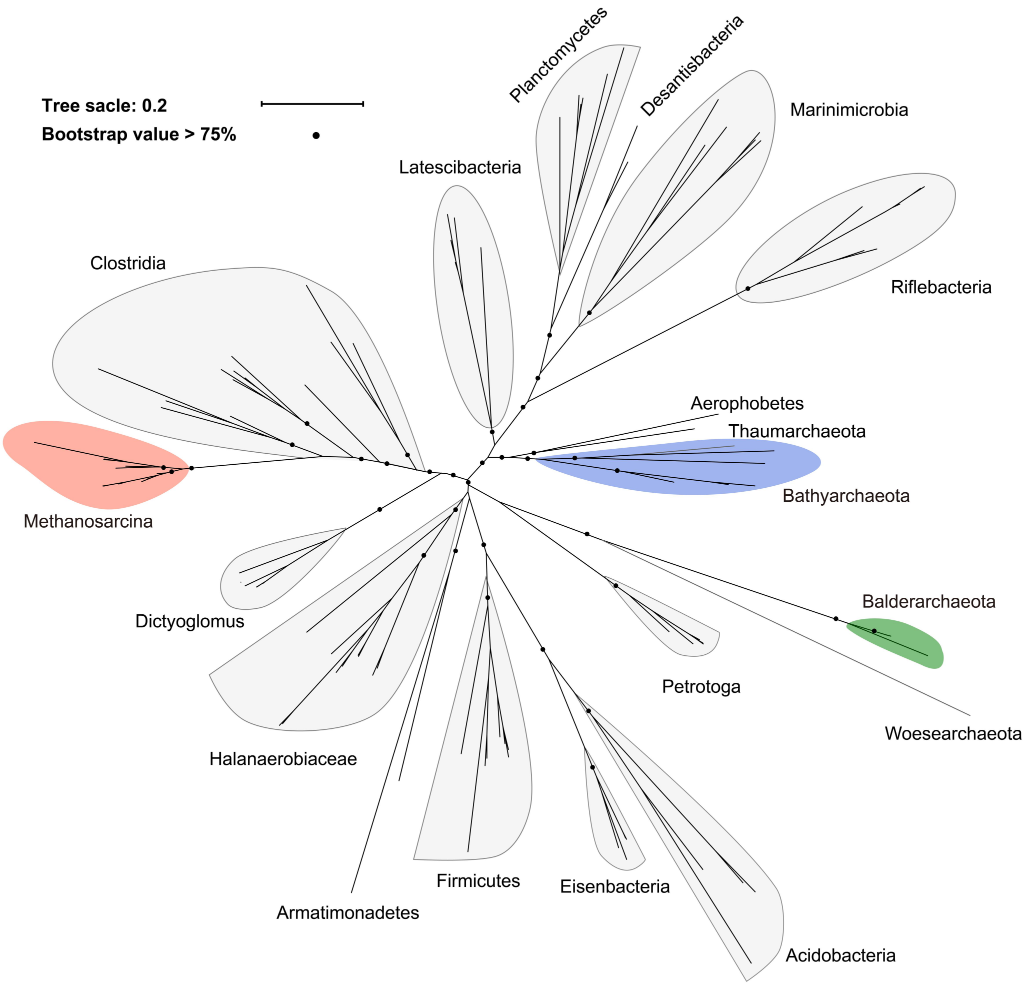


**Supplementary Figure 3** An unrooted maximum-likelihood tree of the Ack homologs using IQ-TREE with best fit model LG+I+G4. Bathyarchaeota in blue, Balderarchaeota in green, Methanosarcina in lightsalmon. The Ack homologs were also found in Woesearchaeota and Thaumarchaeota. Woesearchaeota form a clade with Balderarchaeota and Thaumarchaeota are within Bathyarchaeota lineage, indicating that the *ack* gene in Balderarchaeota and Woesearchaeota acquired from a common bacteria lineage and the *ack* gene in Bathyarchaeota and Thaumarchaeota transferred from a common bacteria donor.

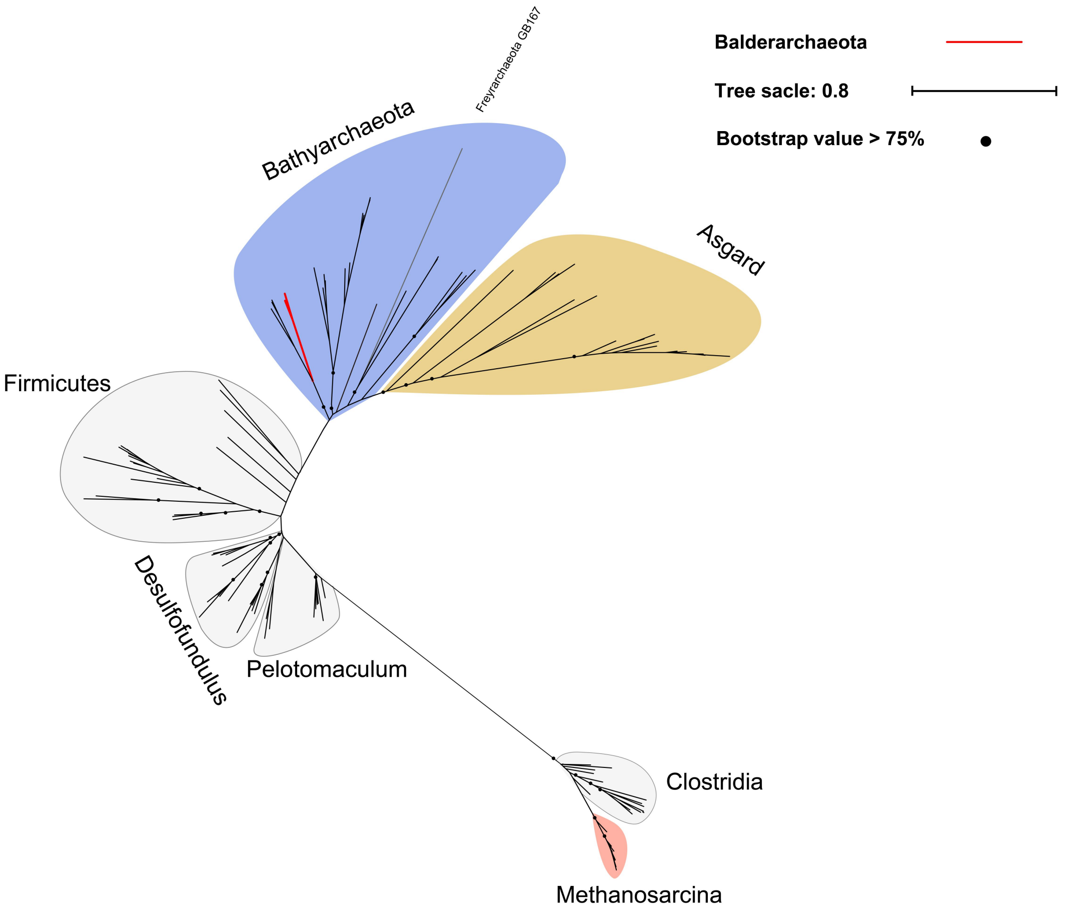


**Supplementary Figure 4** An unrooted maximum-likelihood tree of the Pta homologs using IQ-TREE with best fit model LG+I+G4. Bathyarchaeota in blue, Asgard archaea (except for Balderarchaeota, Hoderarchaeota and Freyrarchaeota) in orange, Methanosarcina in lightsalmon. The red branches are sequences from Balderarchaeota. Both Bathyarchaeota lineage and Asgard lineage are within Firmicutes.
